# *PRDM9* Alleles Guide Distinct Patterns of Recombination Localization in Atlantic Salmon

**DOI:** 10.64898/2026.09.03.749262

**Authors:** Øyvind Sætren Gulbrandsen, Cathrine Brekke, Marie-Odile Baudement, Tan Thi Nguyen, Matthew Peter Kent, Nicola Jane Barson, Sigbjørn Lien

## Abstract

Meiotic recombination in many vertebrate lineages is directed to genomic regions by the DNA-binding protein PRDM9. To date, *PRDM9* research has focused primarily on mammals, and its role in recombination localization has only recently been established in salmonids. PRDM9 binding specificity is determined by a tandemly repeated zinc-finger array that exhibits extensive allelic diversity within and between species. However, the extent to which *PRDM9* variation shapes recombination landscapes in salmonids remains poorly understood. Here, we performed large-scale genotyping of *PRDM9* in an aquaculture population of Atlantic salmon (*Salmo salar*) using long-read sequencing. By integrating *PRDM9* genotypes with pedigree information and genome-wide recombination data, we generated *PRDM9*-specific linkage maps and sets of inferred crossover events. We show that the presence of specific *PRDM9* alleles is associated with differences in recombination localization and identify patterns consistent with dominance interactions between alleles. Using population-averaged recombination rate estimates, we further link a previously described sequence motif enriched in European recombination hotspots to the presence of a common *PRDM9* zinc-finger array allele (RPT6a). Together, our results demonstrate functional differences between *PRDM9* alleles which contribute to divergence in recombination landscapes in Atlantic salmon, paralleling patterns observed in mammals. The broad geographic distribution of Atlantic salmon, combined with extensive pedigree data generated through aquaculture, highlights its potential as a powerful model for studying PRDM9 function and evolution beyond mammals.

## Introduction

Meiotic recombination is a fundamental process in sexually reproducing eukaryotes that ensures proper chromosome segregation during meiosis and reshapes patterns of genetic linkage and genome structure (Hunter, 2015). In many animal species, recombination events are concentrated in short regions known as recombination hotspots, whose locations are largely determined by the activity of the recombination regulator PRDM9 (Baudat et al., 2010; Parvanov et al., 2010; Ségurel et al., 2011). As most research on PRDM9 has focused on mammals, particularly humans and mice, our understanding of its importance, function and diversity in more distant lineages, such as fishes, remains limited.

PRDM9 functions by binding genomic DNA through an array of tandemly repeated C2H2 zinc-finger (ZnF) repeats, which vary in three main aspects: the number of repeats, the DNA-binding affinity of each repeat, and their sequential arrangement. Upon binding, PRDM9 deposits epigenetic marks on nearby histones, thereby facilitating the recruitment of proteins that induce double-stranded breaks and promote crossover events (reviewed in Grey et al., 2018; Paigen & Petkov, 2018). *PRDM9* genes have been identified across vertebrates (Cavassim et al., 2022) and even in insects (Everitt et al., 2025), suggesting an ancient origin and deep evolutionary conservation of PRDM9-directed recombination. Expanding our understanding of *PRDM9*’s function and evolution requires study across distantly related species, as much of our current knowledge is based on mammals and may not be broadly applicable to other lineages. Furthermore, the *PRDM9* gene is remarkably diverse, with allelic variation influencing hotspot localization through differences in the structure and DNA-binding preferences of its ZnF arrays (Berg et al., 2011; Billings et al., 2013; Patel et al., 2016), thereby contributing to substantial variation in recombination patterns across species, populations, and individuals (Marín-García et al., 2024; Pratto et al., 2014; Stevison et al., 2016; Zhou et al., 2018).

Recently, a role of PRDM9 in directing recombination, along with examples of its polymorphism, was demonstrated in Atlantic salmon (*Salmo salar*) and rainbow trout (*Oncorhynchus mykiss*), reinforcing the notion of the widespread function of PRDM9-directed recombination in vertebrates (Raynaud et al., 2025). However, the relationship between specific *PRDM9* variants and recombination hotspot localization has not yet been investigated in Atlantic salmon. Understanding this relationship requires examining how *PRDM9* allelic diversity varies across and within populations, and whether variation in its PRDM9 ZnF array influences the localization of recombination events. Although *PRDM9* variation is expected to shape recombination patterns as in other species, the specific ZnF array features underlying functional differences in DNA-binding activity remain unclear.

PRDM9 has been shown to form multimers where only one of the ZnF arrays contact DNA (Schwarz et al., 2019), enabling one variant to partially, or fully, physically suppress the binding of another (C. L. Baker, Petkova, et al., 2015). This forms a potential mechanistic basis for dominance interactions among *PRDM9* alleles, a phenomenon previously documented in both humans (Pratto et al., 2014; Ségurel et al., 2011) and mice (C. L. Baker, Kajita, et al., 2015; C. L. Baker, Petkova, et al., 2015; Brick et al., 2012). By analyzing recombination patterns across homozygous and heterozygous genotypes, it is possible to simultaneously map hotspot variation and infer potential dominance interactions among PRDM9 alleles. Elucidating these interactions across species could offer critical insight into the molecular basis of dominance in recombination control, which consequently may facilitate better predictions of recombination localization based on *a priori* knowledge about such interactions and has implications for the evolutionary dynamics of *PRDM9*.

Despite its biological importance, the *PRDM9* ZnF array has been difficult to study, largely because its highly repetitive structure makes it difficult to sequence using short-read sequencing. Sanger sequencing is an alternative approach that can resolve these repeats but is both relatively expensive and labor-intensive which limits the scope of *PRDM9* diversity studies. Recent advances in long-read sequencing technologies, including Oxford Nanopore and PacBio platforms, now allow targeted, cost-effective, scalable sequencing of complex genomic regions, enabling accurate genotyping of the complete, hypervariable *PRDM9* ZnF array (Alleva et al., 2021). Although this approach has only been utilized in humans so far (Alleva et al., 2021), it has potential to reveal the extent of *PRDM9* diversity across species, even in non-model species with little-to-no prior knowledge. To establish relationships between *PRDM9* genotypes and recombination outcomes requires extensive recombination data, ideally spanning multiple generations with well-documented pedigrees. Such datasets are rare, but aquaculture populations of Atlantic salmon provide an exceptional resource, offering detailed multi-generational pedigree information and high-density genotyping data. In this study, we leveraged family-based linkage data in combination with large-scale long-read discovery and sequencing of *PRDM9* alleles in the same families to investigate whether allelic variation differentially shapes recombination patterns in Atlantic salmon.

## Methods

An overview of the datasets, derived resources, and analytical workflow used in this study is provided in Figure S6. Briefly, Nanopore-derived *PRDM9* genotypes, SNP array data (Brekke et al., 2023), whole-genome sequencing, recombination hotspots (Gulbrandsen et al., in prep b), and *in silico* motif predictions were integrated to investigate the relationship between *PRDM9* variation and recombination patterns.

### *PRDM9* amplification, pooling, and clean-up

To characterize the extent of variation of the rapidly evolving PRDM9 ZnF arrays, we PCR amplified, and long-read sequenced the *PRDM9* gene from 1213 Norwegian aquaculture Atlantic salmon (*Salmo salar*, n = 1170 excluding repeat samples) from the AquaGen AS breeding program, prioritizing fish with the most available data in Brekke et al. (2023). We used dual barcoding to multiplex and sequence *PRDM9* amplicons. The forward and reverse primers from Raynaud et al. (2025) were extended at their 5’ ends to include 24 extra nucleotides and creating 8 variations for the forward primer and 12 for the reverse (see Table S1). PCR was conducted in 96-well plates where each well received 20 ng genomic DNA, a unique combination of forward and reverse primer (0.5 µM each), LongAmp Hot Start *Taq* 2X Master Mix (New England Biolabs), and nuclease-free water, bringing the final volume to 25 μl. The cycling conditions were: 95°C for 3 min, 35 cycles of 95°C for 30 s, 62°C for 30 s, 72°C for 2 min, a final extension step of 72°C for 7 min, and a hold at 4°C. A negative control (PCR water) was included on each of the PCR plates. After PCR, the concentration of each amplicon was measured using PicoGreen (Molecular Probes, Eugene, OR, USA). Within each plate, the 95 sample amplicons were normalized and pooled to one sample. Each pool was purified using 0.75x AMPure XP beads (Beckman Coulter, Brea, California), and the DNA was eluted in 60 μl nuclease-free water. The concentration of each pool was measured using a Qubit 2.0 dsDNA BR Assay.

### Library preparation and Nanopore sequencing

The pooled sample from each plate was given a second barcode and prepared for ONT sequencing using Native Barcode kit 24 V14 (SQK-NBD114.24) following the ‘Native barcoding Kit 24 V14’ nanopore protocol. Briefly, 163 ng amplicons (130 ng for 1 kb amplicons) from each pool were used as input to prepare the end-prep. Equal amounts of end-prepped DNA from each pool were used for native barcode ligation. The barcoded samples were pooled and cleaned with 0.4x AMPure XP beads (Beckman Coulter, Brea, California). Adaptor ligation was performed using the NEBNext Quick Ligation Module (New England Biolabs). After priming the flow cell, 18ng (equal to 20 fmol) of the final prepared library was loaded into an R10.4.1 flow cell. Sequencing was performed on a PromethION24 operated by MinKNOW v23.04.05 at the Centre for Integrative Genomics (CIGENE), Norwegian University of Life Science (NMBU). Basecalling and initial filtering was done with Guppy v6.5.7 using the “High-accuracy model” and a sequence Q-score cut-off of 9.

### *PRDM9* genotyping

Nanopore output was demultiplexed using Cutadapt v4.2 (Martin, 2011) with an error rate of 0.1, a minimum sequence length of 500 and maximum length of 2500. Demultiplexed sequences were genotyped with a modified version of the human *PRDM9* genotyping pipeline from Alleva et al. (2021), swapping human *PRDM9* ZnFs and flanking query sequences with ones from the Atlantic salmon reference genome (Table S2; Ssal v3.1, GenBank accession: GCA_905237065.2). In samples with sequenced ZnF arrays of multiple lengths, we observed a disproportionate number of sequences corresponding to the shortest array. This bias likely arose from increased PCR amplification of shorter sequences, a phenomenon previously reported by Alleva et al. (2021). To account for the biased recovery of shorter *PRDM9* alleles, we modified the sequence length evaluation thresholds in the genotyper to be more permissive of reporting different-length ZnF arrays (Table S3, Figure S5). This was achieved by reducing the required count ratio of the second most frequent ZnF array length (secondary peak) relative to the most frequent ZnF array length (primary peak). To minimize the risk of this increased permissiveness leading to low-frequency sequencing errors being reported, we increased the required ratio of secondary to tertiary peaks. The modified thresholds were validated by comparing genotyping results between duplicates in this study and with samples also Sanger sequenced in Gulbrandsen et al. (in prep a)

To ensure sufficient sequencing of longer alleles with our lowered threshold for secondary peak detection, we only accepted genotypes based on a minimum of 1000 amplicon sequences containing both intact sequences flanking the ZnF array and tandemly repeated ZnFs. For each sample, no further sequences were processed when a maximum of 2000 sequences fulfilled the above criteria. Among the 32 samples sequenced in duplicate runs, where both runs successfully passed genotyping, we observed a discrepancy in one called allele for a single sample. This sample was excluded from further analysis. Additionally, samples with more than two reported alleles were discarded. The final dataset consisted of 1045 genotyped aquaculture Atlantic salmon.

### *PRDM9* allele nomenclature

To streamline references to *PRDM9* variation in Atlantic salmon (*Salmo salar*), we established a new shorthand naming convention for *PRDM9* alleles. Alleles are named based on: (i) the DNA-binding residues of the second ZnF repeat (IUPAC nomenclature), (ii) the number of ZnF repeats in the array, and (iii) their relative abundance across North American and European wild Atlantic salmon populations (as determined from sequencing in Gulbrandsen *et al*., in prep b). For example, the most frequent *PRDM9* allele in Atlantic salmon aquaculture is named RPT6a. In this case, “RPT” refers to the three amino acid residues at positions –1, +3 and +6 relative to the start of the α-helix domain in the second ZnF, “6” refers to the number of ZnFs in the array, and “a” denotes it is the most abundant allele with the “RPT6” designation. Less frequent alleles starting with “RPT6” are named “RPT6b”, “RPT6c”, and so on. Note that alleles are defined at the nucleotide level, rather than from amino acid residues or only DNA-interacting residues.

### Linkage estimation by *PRDM9* genotype

To test the impact of *PRDM9* genotypes on recombination patterns, we constructed separate linkage maps for families sharing the same *PRDM9* genotype for the most frequent alleles in an Atlantic salmon aquaculture breeding population from AquaGen AS. Sex- and genotype-specific linkage maps were constructed in LepMap3 (Rastas, 2017) using the 35K SNP dataset from Brekke et al. (2023) in the following way: all full-sib families where the father had the same *PRDM9* genotype were used to construct male linkage map for each genotype, and all families where the mother had the same genotype were used to construct female linkage maps. Because alleles QHK5a and QHK5b are identical in the DNA-binding residues of their ZnF arrays, they were treated as the same when constructing linkage maps (QHK5a-b). We constructed linkage maps for each homozygous and heterozygous combination of the three most common functional *PRDM9* alleles (RPT6a, RHT10a, QHK5a-b), yielding six genotype-specific linkage maps.

Physical SNP positions were determined from the Atlantic salmon reference genome, and linkage was estimated within chromosomes. Markers were filtered based on segregation distortion using the *filtering2* module with ‘datatolerance’ set to 0.01, as recommended for multi-family datasets. The *separatechromosomes2* module was run within chromosome, and markers that were not assigned to the main linkage group (LOD < 5) were excluded. To ensure consistent marker order across linkage maps, the *evaluatorder2* module was run with the option to keep marker order fixed when estimating genetic distance (cM). We used the Morgan mapping function, as recommended for medium- to high-density marker data where double crossovers between markers are unlikely (Kivikoski et al., 2023).

To avoid recombination rate inflation caused by genotyping errors, SNPs spaced less than 500 bp apart were excluded. Only SNPs shared among all six linkage maps were retained to ensure comparable information density. In Atlantic salmon, male recombination is largely confined to repeat-rich telomeric regions, whereas female recombination is more evenly distributed along the genome (Brekke et al., 2023), a pattern that we confirmed also applies to the genotype-specific linkage maps constructed here. All downstream analyses therefore focused on female recombination, which allows more effective identification and comparison of *PRDM9*-specific recombination patterns across less repetitive genomic regions.

To restrict analyses to regions of high female recombination, we used a previously published female-specific linkage map constructed from 5568 unique full-sib families from the same breeding population (Brekke et al., 2023). For each chromosome, high-recombination regions were defined using a 1 Mb– averaged map as the longest consecutive interval on each chromosomal arm where recombination rate (cM/Mb) exceeded either the chromosome-wide mean or 1 cM/Mb, whichever was lower. Up to two consecutive windows below this threshold were permitted within each interval. Regions outside these intervals were excluded (Figure S1). After filtering, 14 737 SNPs remained in each linkage map.

To infer the resolution at which we could compare linkage maps while retaining information for compared regions, we considered SNP density across multiple window sizes. For each of the six *PRDM9*-genotype-specific linkage maps, recombination rates were averaged for eleven window sizes ranging from 50 kb to 1 Mb. As window size decreased, an increasingly larger proportion of windows did not contain any SNPs. To balance the information content of windows with resolution we opted for a window size of 200 kb, corresponding to 9.1% of windows with no SNPs and 76.7% windows with two or more SNPs (Figure S2). The final set consisted of 4160 windows to be compared between the six

*PRDM9*-specific linkage maps.

### Whole genome sequencing and variant calling

Whole genome resequencing (WGS) was performed on 443 Atlantic salmon individuals originating from AquaGen AS breeding populations, including both historical founders and more recent year classes.

Genomic DNA was extracted using standard protocols and sequenced across multiple platforms, including BGISEQ-500 and Illumina, to ensure broad coverage and representation. Raw sequencing reads were aligned to the Atlantic salmon reference genome assembly Ssal_v3.1 (GenBank accession: GCA_905237065.2) using BWA-mem2 (Vasimuddin et al., 2019), with PCR duplicates identified and fl(Faust & Hall, 2014)ER v0.1.26 (Faust & Hall, 2014). To maintain consistency across data derived from different sequencing technologies, a customized pipeline incorporating SAMtools (Danecek et al., 2021) and GATK4’s *BaseRecalibratorSpark* and *ApplyBQSRSpark* was employed for base quality score recalibration (Van der Auwera & O’Connor, 2020). Variant discovery was carried out using the DeepVariant v1.1.2 pipeline in whole-genome mode, and resulting calls were merged across samples using GLNEXUS v1.4.1 (Yun et al., 2021). Filtering followed established benchmarks, emphasizing heterozygosity excess over allele balance, and resulted in the retention of over 16 million high-confidence bi-allelic SNPs suitable for downstream analyses.

### Crossover inference from whole genome sequenced families

To identify recurring binding motifs at crossover locations in Atlantic salmon, we used WGS data to detect crossover events in full-sib families with known parental genotypes. Like we did for the linkage maps, we focused on female recombination events. From a total of 443 whole-genome–sequenced Atlantic salmon, we selected WGS data for 18 mothers across 21 full-sib families. Each family included the fathers and 2-7 offspring, enabling phasing of offspring gametes and fine-scale crossover event detection for the female meioses. Quality control of the WGS data was conducted with PLINK v1.9 (Chang et al., 2015). Called SNPs were filtered to remove markers with minor allele frequency lower than 0.01 and markers deviating from Hardy-Weinberg equilibrium (exact test p-value < 0.001, using the mid-p adjustment). Finally, markers and samples with a Mendel error rate exceeding 0.01 were removed (no samples were lost due to Mendelian errors). The final dataset comprised 3 719 113 segregating SNPs, with an average genome-wide density of ∼13 SNPs per 10 kb. Linkage mapping was done with LepMap3 with the same parameter settings as for the *PRDM9* genotype-specific linkage maps. Individual crossover positions were detected from the phased gamete output from the LepMap3 *evaluateorder2* module. Only crossovers originating from female meioses were retained for downstream analyses.

### *PRDM9* allele imputation in crossover dataset

Only four of the individuals in the WGS dataset used to infer crossovers had been *PRDM9* genotyped, preventing us from categorizing crossovers by parental *PRDM9* allele presence. To address the incomplete *PRDM9* genotype information for the parents in the crossover families, we imputed the missing genotypes by linking available WGS SNPs and *PRDM9* genotypes from the same individuals. Utilizing WGS from 443 Atlantic salmon, we first determined which SNPs were linked to *PRDM9* by estimating the surrounding linkage blocks. As we were working with a closed breeding population, we assumed consistent *PRDM9*-linked haplotypes without introgression. The 15 SNPs overlapping the hypervariable ZnF array-encoding region of PRDM9 were removed to account for unreliable mapping, before phasing chromosome 5 with Beagle v.5.4 (Browning et al., 2021) using default parameters. D’ and LD blocks were estimated in HaploView v.4.2 (Barrett et al., 2005) from 50 kb on either side of the ZnF array-encoding region, skipping the quality filtering step. For the *PRDM9* genotyped individuals, a haplotype was defined from the 57 phased SNPs on the LD block encompassing the region encoding the PRDM9 ZnF array. From the imputation WGS dataset, 53 individuals had available *PRDM9* genotypes, possessing 8 different alleles. Examining haplotype and allele presence in these 53 individuals, we found that 11 out of 14 *PRDM9*-linked haplotypes exclusively co-occurred with a single *PRDM9* allele (Table S4).

Using the 11 haplotypes uniquely linked to *PRDM9* alleles, we imputed *PRDM9* genotypes for 85 out of 99 individuals used for crossover inference, including mothers, fathers, and offspring. For all but one family, which was discarded, all imputed offspring genotypes could be derived by crossing the parental genotypes. Additionally, the imputed genotypes corresponded to *PRDM9* genotypes for the four individuals in the crossover data set that we had *PRDM9* sequenced. We selected datasets for crossover inference from the 17 families where both maternal alleles were successfully imputed, and in which both parents were imputed to be homozygous for either allele RPT6a or QHK5a. These two *PRDM9*-specific crossover sets consisted of 4 families with 14 offspring in total (RPT6a, 2-5 offspring per family), and 2 families with 2 and 5 offspring (QHK5a). Additionally, we analyzed the full crossover set of 21 families where parents were successfully imputed (“ungenotyped”), resulting in 3 sets of crossover regions: ungenotyped, homozygous RPT6a, and homozygous QHK5a.

### *De novo* crossover motif prediction

For the three crossover sets (ungenotyped, homozygous RPT6a, and homozygous QHK5a), inferred crossover regions were removed if they fell within 1 Mb of the chromosome ends or were over 5 kb in length. *De novo* motif prediction was performed independently for the homozygous RPT6a, homozygous QHK5a, and the total set of crossovers (ungenotyped) by XSTREME v.5.5.7 (Grant & Bailey, 2021) in the web suite, with one expected occurrence per sequence, a minimum motif width of 6, and a maximum motif width of either 3 times the number of ZnFs in the array (assuming 3 DNA-interacting residues per ZnF) or 24 nucleotides (example human PRDM9 motif length). A total of 23 unique motifs were identified across all crossover regions (Table 1).

**Table 1.**
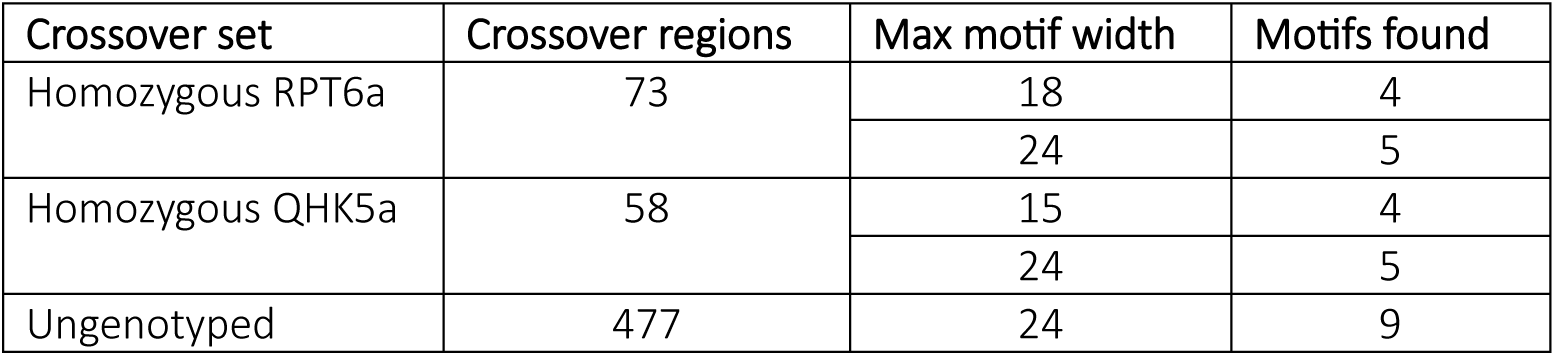
The PRDM9 background of three sets of inferred crossovers used in de novo motif prediction w/ EXTREME. Two different motif widths were run for PRDM9-specific crossover sets (“Max motif width”).

| Crossover set | Crossover regions | Max motif width | Motifs found |
| --- | --- | --- | --- |
| Homozygous RPT6a | 73 | 18 | 4 |
|  |  | 24 | 5 |
| Homozygous QHK5a | 58 | 15 | 4 |
|  |  | 24 | 5 |
| Ungenotyped | 477 | 24 | 9 |

### Hotspot motif enrichment and motif recombination rate

To identify *PRDM9* associated DNA sequence motifs within recombination hotspots, we looked for motif enrichment in wild population recombination hotspot regions and tested for recombination rate increases at motif-matching sites. We assessed motif enrichment in population-specific hotspot regions identified in Gulbrandsen et al. (in prep b), these hotspots were inferred from population-averaged historical recombination estimates from 30 wild Atlantic salmon populations across North America and Europe. Motifs were inferred using several parallel strategies: (i) PRDM9 binding motifs inferred by XSTREME from genotype-specific crossover regions, (ii) *in silico* motif predictions for alleles RPT6a and QHK5a using DeepZF (Aizenshtein-Gazit & Orenstein, 2022) and a polynomial SVM (https://zf.princeton.edu/: Persikov et al., 2009; Persikov & Singh, 2014), and (iii) rainbow trout (*Oncorhynchus mykiss*) motifs inferred from ChIP-Seq and a Atlantic salmon (*Salmo salar)* motif defined from its enrichment in European population hotspots, both from Raynaud et al. (2025). The *PWMpredictor* step of the DeepZF pipeline was run for the full-length ZnF arrays, including the 40 flanking amino acids on either end of each ZnF.

Hotspot sets were screened for enrichment of PRDM9 binding motifs in each population using SEA v.5.5.5 (Bailey & Grant, 2021) with a 6^th^ order Markov (--m 5) background frequency computed from the Atlantic salmon reference genome and a q-value cutoff of 0.05. Control sequences were generated by randomly sampling a corresponding set of genomic regions matching the size, chromosome, and GC content (± 1%) of hotspot regions, disallowing overlap of hotspots or control regions sampled in the same iteration. Control sequence sampling was independently performed 500 times for each population hotspot set. Genomic GC content was estimated by averaging each chromosome over a 2 kb sliding window, moving 100 bp at a time. For each motif enriched in a set of population hotspots, the log-scaled recombination rates in control sequences containing the motif (as reported by SEA) were tested against control sequences not containing the motif for an increase in mean recombination rate using a one-sided Welch’s t-test. Final *p*-values were Bonferroni corrected.

## Results

### *PRDM9* diversity in aquaculture Atlantic salmon

To investigate the diversity of *PRDM9* in a salmonid population, we sequenced and genotyped the ZnF arrays in 1045 aquaculture salmon (AquaGen AS) using Oxford Nanopore long-read technology. The genotyping revealed 14 unique *PRDM9* alleles, with ZnF array lengths ranging from 5 to 10 consecutive ZnF repeats. The four most frequent alleles, RPT6a (52.6%), QHK5a (24.1%), RHT10a (6.7%), and QHK5b (4.4%), accounted for more than 87% of the observed *PRDM9* diversity (Figure 1; see “*PRDM9* allele nomenclature” for naming conventions). Among these, RHT10a encodes a ZnF array consisting of 10 repeat units, twice the length of QHK5a and QHK5b. The most common allele (RPT6a) possesses six ZnF repeats. The remaining 12.2% of *PRDM9* variation was distributed among nine additional alleles, two of which were observed only once, highlighting the presence of rare variants in the population.

**Figure 1.**
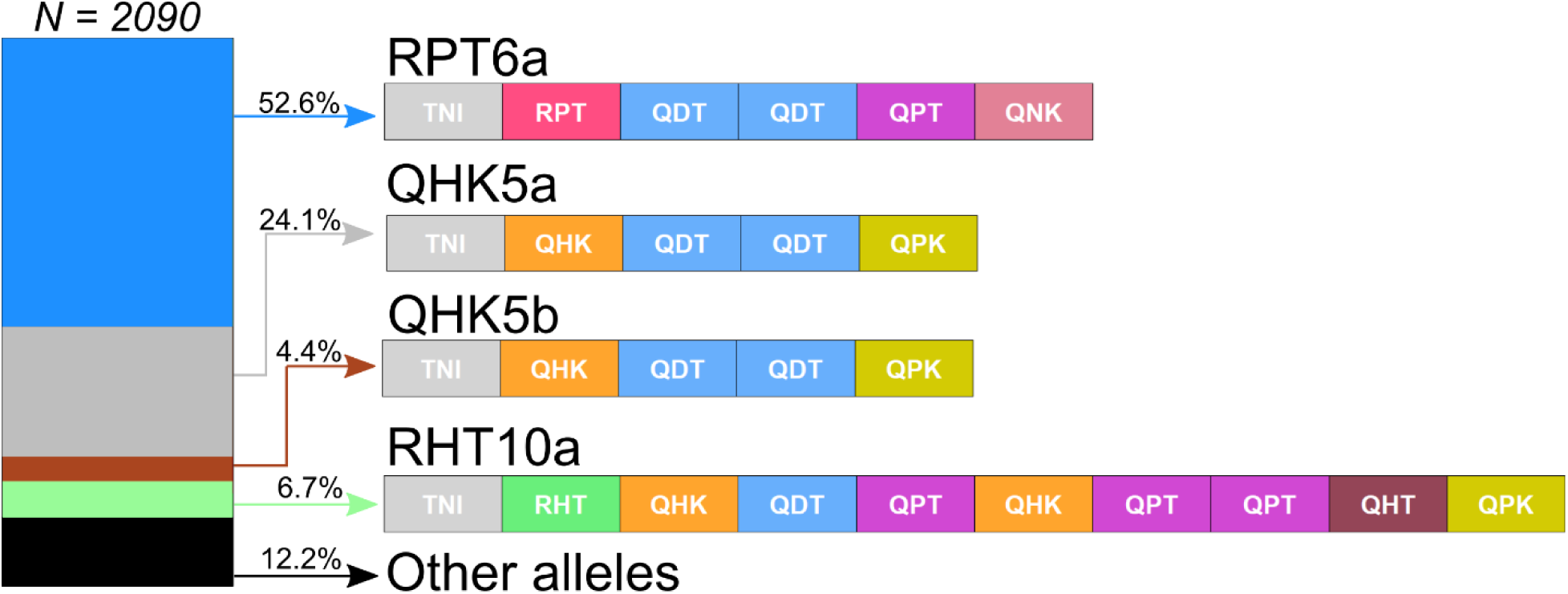
Relative proportions of the four most common PRDM9 alleles (left) and their corresponding ZnF arrays (right), genotyped from 1045 aquaculture Atlantic salmon (Salmo salar). Each ZnF is annotated with the amino acid contact residues at positions −1, +3 and +6 in relation to the start of the α-helix.

From the 2090 genotyped *PRDM9* ZnF arrays, we identified 15 unique 84 bp length ZnF repeat units. To assess potential functional differences, we grouped these into nine putatively functionally distinct repeat units based on the DNA-binding amino acids which determine binding specificity at positions −1, +3, and +6 relative to the start of the α-helix domain. Consistent with PRDM9 in other species, the first ZnF repeat in each array was conserved, containing threonine at position −1, asparagine at +3, and isoleucine at +6. In contrast to the canonical pattern observed in other species, the final ZnF repeat in each array consistently differed in structure from the preceding repeats, notably lacking the terminal zinc-coordinating histidine that normally stabilizes the ZnF folding. This unusual structural feature of the terminal ZnF has been previously noted by Raynaud et al. (2025) and may have implications for DNA-binding specificity at the ends of the array.

### *PRDM9* genotypes differ in recombination enrichment patterns

To test differences between recombination localization across different *PRDM9* genotype backgrounds, we generated six maternal genotype-specific linkage maps. Linkage maps were constructed for groups of families in which the mothers shared one of the four most abundant *PRDM9* alleles in the farmed salmon strain (RPT6a, RHT10a, QHK5a, or QHK5b; Figure 2c). Because the alleles QHK5a and QHK5b have identical DNA contact residues (Figure 1), they were grouped as a single functional variant, QHK5a-b. First, we calculated genome-wide recombination in 200-kb windows and identified consecutive regions of elevated female recombination rates (Figure S1, see “Linkage estimation by *PRDM9* genotype”). Windows were flagged as a recombination peak if they met both of the following criteria: (i) the recombination rate exceeded three times the genomic average, and (ii) flanking windows were below two times the genomic average. To ensure consistent comparisons across all six linkage maps, 200-kb windows lacking a recombination peak in any map were removed, leaving 436 windows for comparisons. We then calculated Jaccard indices for each of the 15 pairwise combinations of PRDM9 genotype specific linkage maps (Figure 2a). Statistical significance was assessed using a two-tailed Fisher’s exact test, using Bonferroni-corrected p-values (Figure 2a).

**Figure 2.**
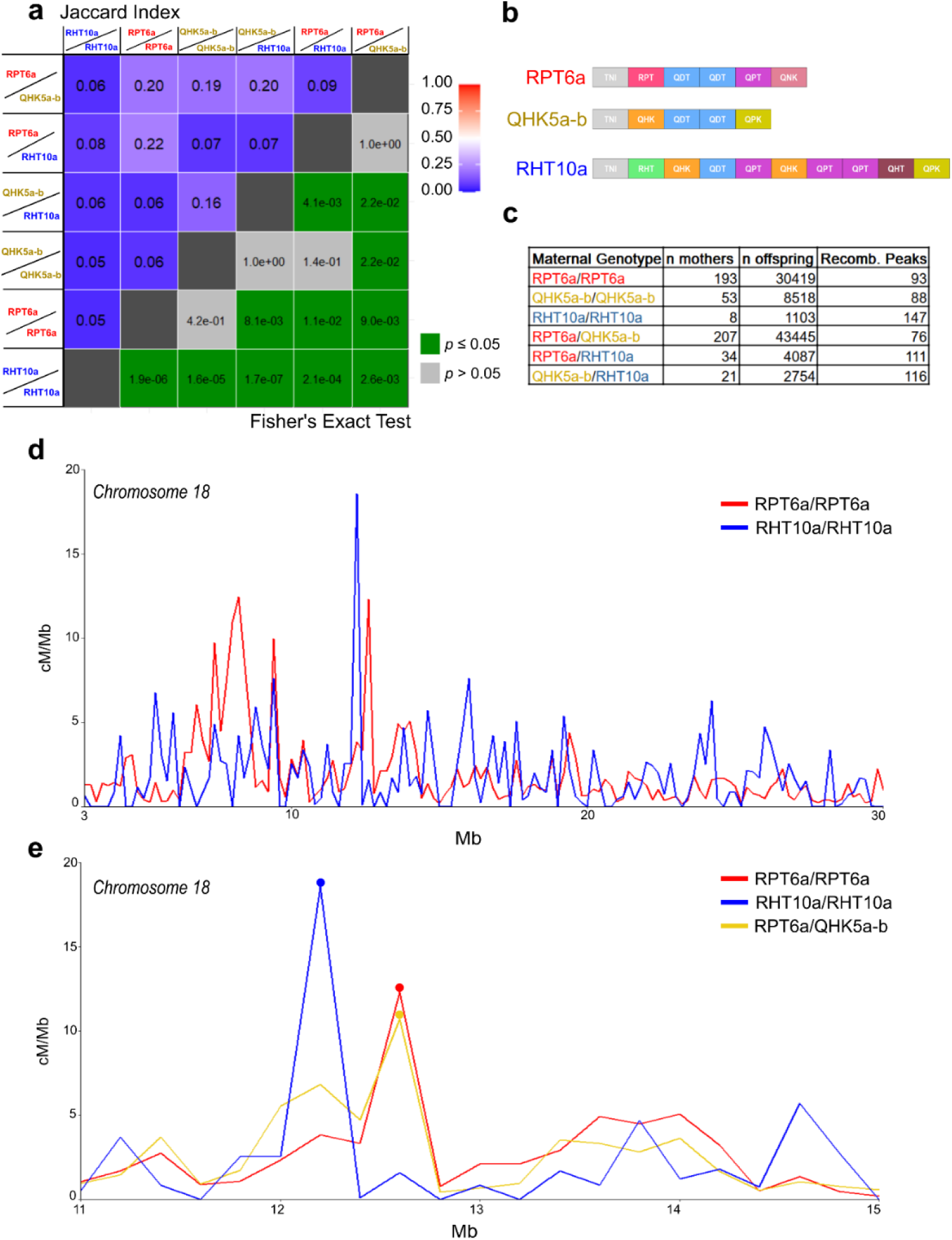
Recombination pattern differences in linkage maps inferred from family sets with different PRDM9 backgrounds. (a) Pairwise comparisons of recombination patterns in linkage maps inferred from 6 different maternal PRDM9 genotypes. Jaccard similarity indices are in the upper left and results of two-tailed Fisher’s exact test after Bonferroni correcting p-values are in the bottom right. Only windows with a recombination peak in at least one genotype were compared. (b) ZnF array structures of the most common PRDM9 alleles in aquaculture Atlantic salmon. Linkage maps were generated from families where mothers possessed combinations of these alleles. Each ZnF is annotated with the encoded amino acid contact residues at positions −1, +3 and +6 in relation to the start of the α-helix. (c) Summary counts of individuals included to generate each linkage map. By PRDM9 genotype: the number of mothers, the total number of offspring derived from the mothers, the number of 200-kb windows identified as recombination peaks. (d) Recombination estimates (1-Mb averaged) for two of the linkage maps along chromosome 18, showing different recombination activity between PRDM9 genotypes. Only the consecutive interval of high female recombination which was used for the analyses is plotted. (e) Recombination estimates (200-kb averaged) for three of the linkage maps along an example interval. Dotted peaks were flagged as recombination peaks and included in comparisons. Here, recombination peaks overlap in two linkage maps which share an RPT6a allele, while the RHT10a/RHT10a recombination peak does not. Note that broad recombination peaks are not selected to ensure comparisons at similar scales and precision.

Jaccard coefficients for recombination hotspot overlap broadly fell into two ranges (Figure 2a): a low overlap range (0.05 to 0.09) and a higher overlap range (0.16 to 0.22). Overlap was generally greater when two genotypes shared an allele (e.g. RPT6a/QHK5a-b vs. RPT6a/RPT6a and RPT6a/QHK5a-b vs. QHK5a-b/RHT10a, Jaccard coefficient = 0.20) compared to genotypes that did not share alleles (e.g. QHK5a-b/RHT10a vs. RPT6a/RPT6a and RPT6a/QHK5a-b vs. RHT10a/RHT10a, Jaccard = 0.06). This pattern supports the hypothesis that different *PRDM9* alleles direct recombination to distinct genomic locations in Atlantic salmon. Interestingly, recombination peak overlaps were consistent across all three comparisons of homozygous *PRDM9* genotypes, indicating that each encoded PRDM9 variant targets distinct, largely non-overlapping genomic sequences. These results suggest that shared alleles partially align recombination patterns between families, whereas differing alleles drive genotype-specific hotspot localization.

Moreover, recombination overlaps suggest a dominance effect of other *PRDM9* alleles over the RHT10a allele. Specifically, recombination peaks in the linkage map from maternal RHT10a/RHT10a genotypes are as different to the heterozygote lacking an RHT10a allele (RPT6a/QHK5a-b) as they are to the two heterozygotes containing the allele (RPT6a/RHT10a and QHK5a-b/RHT10a). The dominance of QHK5a-b over RHT10a is further supported by the similarity in recombination peak overlap between RPT6a/QHK5a-b and QHK5a-b/RHT10a (Jaccard = 0.20, p = 0.022), which is comparable to the overlap observed when RHT10a is replaced by QHK5a-b (e.g. RPT6a/QHK5a-b vs. QHK5a-b/QHK5a-b, Jaccard = 0.19, p = 0.022). These results suggest that RHT10a has a reduced influence on recombination localization in the presence of QHK5a-b. In another heterozygote-heterozygote comparison we find a low recombination overlap, consistent with the effect of RHT10a suppression in one or both genotype backgrounds (RPT6a/RHT10a vs. QHK5a-b/RHT10a, Jaccard = 0.07, p = 0.0041), further supporting a dominance interaction. However, we do not observe a dominance effect in the final heterozygote-heterozygote comparison (RPT6a/QHK5a-b vs. RPT6a/RHT10a, Jaccard = 0.09, p = 1.00), even though a similar Jaccard index to RPT6a/QHK5a-b vs. QHK5a-b/RHT10a would be expected if both QHK5a-b and RPT6a are dominant over RHT10a. This discrepancy suggests that the dominance effect may be stronger for the QHK5a-b alleles, or that the number of recombination peaks in the RPT6a/QHK5a-b vs. RPT6a/RHT10a comparison was insufficient to detect such an effect.

We note that 4 out of 15 comparisons were statistically insignificant (p > 0.05). This lack of significance appears to result from fewer recombination peaks in certain comparisons, particularly those involving maternal genotypes QHK5a-b/QHK5a-b and RPT6a/QHK5a-b, which also have the lowest total number of recombination peaks (Figure 2c). Despite the lack of statistical significance for some comparisons, the overall trend remains clear: recombination peak overlap is greater when maternal genotypes share *PRDM9* alleles and lower when they do not.

### PRDM9 motif enrichment in recombination hotspots

Since *PRDM9* genotypes appear to influence recombination localization in Atlantic salmon, we next aimed to identify genomic sequences that are overrepresented and specifically associated with different *PRDM9* alleles. Using recombination hotspots inferred from population-level recombination maps of wild Atlantic salmon across Europe and North America Gulbrandsen et al. (in prep b), we tested a panel of candidate PRDM9 binding motifs for enrichment within hotspot regions. These motifs were derived independently of the population-level recombination maps and originated from multiple sources, including motifs inferred from fine-scale crossover regions identified using whole-genome sequencing in families where the mother was homozygous for the *PRDM9* alleles RPT6a or QHK5a, in silico binding predictions for the same alleles, and previously described salmonid PRDM9-associated motifs (Raynaud et al., 2025). The recombination hotspots represent historic, population-averaged recombination, integrating recombination across sexes and across all *PRDM9* alleles that have contributed to recombination in each population. To test candidate PRDM9 binding motifs for allele-associated enrichment, we conducted a motif enrichment analysis using SEA (Bailey & Grant, 2021), testing 30 candidate motifs for association with recombination hotspots across 30 wild populations, 16 of which also had available *PRDM9* genotype data (Figure 3 and Figure S3). To further refine candidate motifs, we tested whether hotspot-enriched motifs were associated with increased log-scaled recombination rates compared to control sequences lacking the motif.

**Figure 3.**
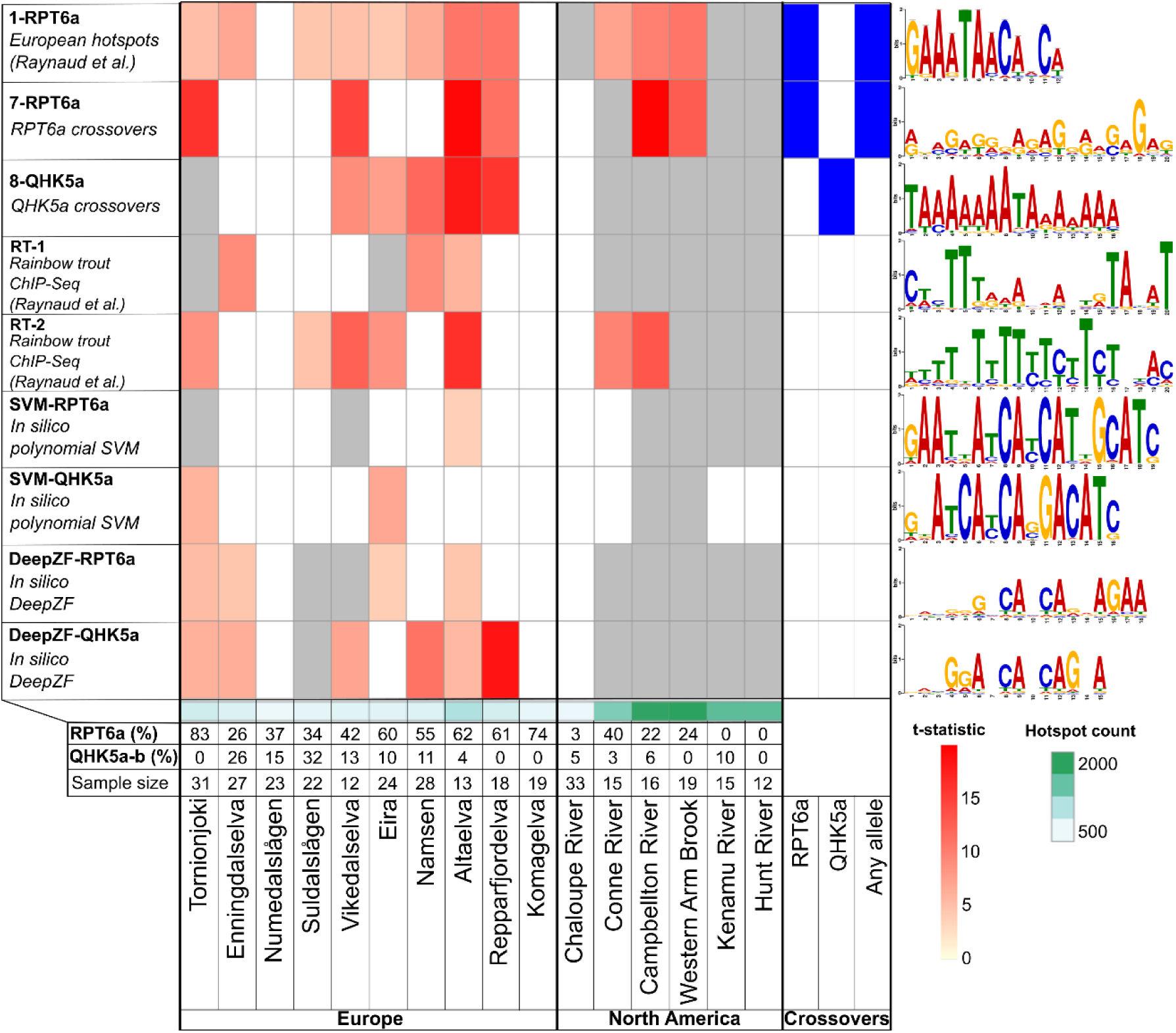
Selected motifs significantly enriched in hotspot regions and their associated increases in recombination intensity in genomic regions containing a PRDM9 binding motif candidate across wild Atlantic salmon populations. The heatmap shows the results of SEA enrichment analyses of hotspot regions, as well as one-sided Welch’s t-tests for recombination rate increases in comparable genomic regions with and without a PRDM9 binding motif candidate (left panel: motif names, origin and prediction method, right panel: corresponding sequence motif logos), across 16 wild Atlantic salmon populations (bottom) with PRDM9 genotypes and recombination estimates generated in Gulbrandsen et al. (in prep b). The number of individuals genotyped for PRDM9 and proportions of relevant alleles are listed for each population. Motifs not significantly enriched in the hotspot set of a population (white) were excluded from recombination rate tests. The increase in recombination at motif-matching sites (t-statistic, yellow to red) is shown for motifs with significant recombination rate increases (Bonferroni-corrected p ≤ 0.05). Motifs significantly enriched in hotspots but without significant recombination rate increases are shown in grey. Crossover regions were not tested for recombination rate increases, but motifs enriched within crossover regions are colored blue. The number of hotspots identified for each population is at the bottom row of the heatmap (white to dark green). For complete results, including populations without genotypes and all tested binding motif candidates, see Figure S3.

Overall, we did not find clear patterns of significantly enriched motifs associated with either phylogeographical group or specific *PRDM9* alleles. This outcome may reflect the combined influence of multiple *PRDM9* alleles shaping historical recombination, as well as potential imprecision in the candidate binding motifs. Notably, PRDM9 binding motifs inferred from rainbow trout ChIP-seq data (Raynaud et al., 2025) were enriched in hotspots to a degree comparable to several other candidate motifs and showed elevated recombination in a subset of populations (Figure 3 and Figure S3).

Additionally, higher recombination rates were found at motifs resembling dinucleotide repeats (highlighted in red, Figure S3), an association which has been found in other species with and without PRDM9-directed recombination (Bagshaw et al., 2008; Majewski & Ott, 2000). We also observed a trend toward a higher number of significantly enriched motifs in populations with more recombination hotspots, a pattern that did not correspond to the distribution of any specific *PRDM9* allele. Enriched motifs within crossover regions generally reflected their source sequences, and no shared enriched motifs were detected between crossover regions of *PRDM9* alleles RPT6a and QHK5a. These findings suggest that the high number of motifs and lack of consistent patterns across populations likely arise from variability in underlying recombination estimates and differences in the sequence composition of population specific hotspots.

The motif identified by Raynaud et al. (2025) as overrepresented in European recombination hotspots (*1-RPT6a*) was also the motif most frequently enriched in wild Atlantic salmon population hotspots in our analysis. Specifically, *1-RPT6a* is enriched in 23 out of the 30 populations, across all four phylogeographic groups (Figure S3). In most populations, *1-RPT6a* exhibits the highest enrichment ratio, calculated as the proportion of motif matches in hotspots divided by proportion of matches in control sequences (Figure S4), and consistently ranks near the top in the SEA enrichment analysis. It is possible that *1-RPT6a* represents a composite PRDM9 binding motif, as it was derived from recombination hotspots across diverse European populations with a currently wide range of different alleles and unknown allele history, yet it appears strongly biased towards the sequence target of most common European allele, RPT6a. Supporting this, the SEA analysis identified *1-RPT6a* as the second-ranking motif in RPT6a specific crossovers, while a similar motif ranks higher (*2-AATAAAAAAAAGAG*; Figure S3). However, this latter motif was generated from the same sequences it was tested against, likely introducing bias favoring it over the Raynaud motif. Finally, *1-RPT6a* also shows a high enrichment ratio in ungenotyped crossovers (Figure S4), which are likely predominantly influenced by the RPT6a allele due to its prevalence in Atlantic salmon aquaculture (Figure 1).

The *1-RPT6a* motif showed a significant, albeit modest, increase in recombination rate at matching sites in most populations (Figure 3 and Figure S3). Notably, although *1-RPT6a* was enriched in hotpots of many North American populations, significant recombination rate increases at motif-matching sites were observed only in the three North American populations with moderate presence of RPT6a alleles. These recombination rate increases, combined with high enrichment ratios (Figure S4) and widespread enrichment across populations, collectively identify *1-RPT6a* as the strongest candidate for *RPT6a* binding motif. We also propose *7-RPT6a* (consensus sequence: RDMGAGRGAGAGGRAGAGAG) as an alternate binding motif candidate, as it displays similar enrichment patterns and recombination rate increases as *1-RPT6a*. Given the considerable sequence differences between the two motifs, *7-RPT6a* likely represents a distinct motif rather than a derivate arising from limited data.

We did not find consistent enrichment patterns for QHK5a motifs, which is unsurprising given their low frequency and even distribution across most populations. Additionally, the lower number of QHK5a crossovers compared to RPT6a crossovers likely reduced the precision of motif recovery from these sequences. Nevertheless, we propose *8-QHK5a* (consensus sequence: TAAAAAAATAAAAAAA) as our best candidate for a QHK5a binding motif based on our current data (Figure 3). However, its reliability is uncertain, as *8-QHK5a* is enriched in four populations where QHK5a, or its likely functional equivalent QHK5b, is absent (Tornionjoki, Repparfjordelva, Western Arm Brook, Hunt River). Conversely, the motif is not enriched in two populations with a high proportion of these two alleles (Enningdalselva, Suldalslågen). Nonetheless, a significant increase in recombination activity at *8-QHK5a* sequence sites in a geographically close cluster of European populations carrying these alleles supports the notion that the motif may represent historical recombination sites.

Enrichment and recombination rate increases for *in silico* predicted binding motifs indicate that DeepZF-predicted motifs align more closely with regions of elevated recombination than those generated by polynomial SVM. This is supported by DeepZF motif enrichment across more populations and higher recombination rates at motif-containing sites (Figure 3 and Figure S3). However, DeepZF motif predictions for the RPT6a and QHK5a alleles tend to overlap in both enrichment and higher recombination, regardless of the underlying population *PRDM9* genotypes in each population. It remains unclear whether the overlap of predicted motifs reflects an inability of DeepZF to distinguish between RPT6a and QHK5a specifically, or from a general limitation in distinguishing salmonid *PRDM9* alleles.

## Discussion

In this study, we demonstrate that variation in the *PRDM9* ZnF array in aquaculture Atlantic salmon is associated with distinct recombination patterns. Using long-read Nanopore sequencing and a modified version of the human *PRDM9* genotyping pipeline developed in Alleva et al. (2021), we successfully genotyped *PRDM9* in 1045 Atlantic salmon and identified 14 alleles, four of which were predominant in the aquaculture population (RPT6a, QHK5a, RHT10a, and QHK5b). By integrating these genotypes with multi-generational pedigree data, we inferred regions of elevated recombination associated with distinct *PRDM9* genotype backgrounds. Different *PRDM9* alleles were associated with distinct recombination landscapes, with heterozygous individuals generally exhibiting intermediate recombination patterns relative to the corresponding homozygotes. A notable exception was the RHT10a allele, whose effect on recombination localization was largely suppressed in heterozygotes, suggesting dominance of other alleles over RHT10a in shaping recombination placement.

Extending these analyses to population-level recombination landscapes, we screened a diverse panel of putative PRDM9 binding motifs against recombination hotspots inferred from wild Atlantic salmon populations across Europe and North America. This analysis revealed a strong association between a previously described motif enriched under European hotspots (Raynaud et al., 2025) and the presence of the RPT6a allele in European and some North American populations. Beyond this signal, we found limited or no evidence for candidate motifs associated with the other tested allele, QHK5a, or for alternative RPT6a motifs derived from crossover-based inference or in silico binding predictions. This likely reflects both constraints of the crossover datasets used to infer allele-specific motifs and the limited precision of current in silico binding prediction approaches.

### Long-read sequencing of *PRDM9* facilitates high-resolution populational surveys across species

The tandemly repeated structure encoding the *PRDM9* ZnF array presents a fundamental obstacle to accurate genotyping, as it requires both determination of array length and resolution of the SNPs encoding DNA-contacting residues in each repeat. High sequence similarity among repeats, coupled with frequent tandem duplications, prevents reliable assembly from short-read sequencing data. In most studies to date, Sanger sequencing has been the preferred tool to generate accurate sequences spanning the length of the ZnF array without the need for assembly (Baudat et al., 2010; Raynaud et al., 2025). However, Sanger sequencing is costly and does not scale well, a problem further exacerbated by the need for multiple rounds of sequencing to distinguish potential heterozygous genotypes when alleles are of equal length.

Long-read sequencing technologies, such as PacBio and Oxford Nanopore, overcome the limitations of short-read sequencing by spanning the entire ZnF array in a single read, bypassing assembly of a repetitive region and keeping haplotypes separate. Although base-accuracy from long-read technologies can be lower than from Sanger sequencing, this difference is becoming less significant thanks to improvements in long-read chemistry and software and, when combined with multi-pass sequencing and/or high-coverage, large-scale and cost-effective long-read based genotyping of thousands of samples is achievable. The *PRDM9* genotyping pipeline developed by Alleva et al. (2021) provides a verified means of leveraging long-read data for genotyping, and our study demonstrates that this approach can be extended beyond humans.

When applying this genotyping method to Atlantic salmon, we recovered diploid genotypes for 89.2% of individuals which is slightly lower than the 95.7% success rate achieved in humans by Alleva et al. (2021). This difference likely reflects lower duplicate sequencing depth and more stringent coverage thresholds applied here, rather than limitations in transferability. Adapting the tool for use in salmon required only minor modifications, such as replacing reference sequences to the species of study. However, to address potential biases in PCR amplification favoring shorter ZnF arrays when allele lengths differ, we recommend adjusting ZnF array length evaluation thresholds to align with the input sequence length distribution. Importantly, this adjustment can be made post-sequencing and does not require prior knowledge of *PRDM9* variation.

Our successful application of this *PRDM9* genotyping tool to Atlantic salmon, a species with a greater range of *PRDM9* variation than humans, highlights its potential for diversity screening across species. Scalable, long-read-based *PRDM9* genotyping enables high-resolution analyses that would be infeasible with traditional sequencing approaches. As such, this framework opens the door to a better understanding of the structure of *PRDM9* diversity and comparative studies of *PRDM9* evolution and function across taxa with PRDM9-directed recombination.

### Sequence and functional diversity among *PRDM9* alleles

Of the 14 *PRDM9* alleles detected in the aquaculture population sequenced in this study, the alleles RPT6a and the synonymous variants QHK5a/QHK5b (QHK5a-b) were notably more frequent, at 52.6% and 28.5%, respectively. *PRDM9* diversity in humans and mice spans a wide range, from near monomorphism with a 90% frequency of human allele A in a Finnish population (Alleva et al., 2021) to the coexistence of multiple alleles at comparable frequencies in several populations of Madeira house mice (Vara et al., 2019). The Atlantic salmon population examined here conforms to a commonly observed pattern in which one or two alleles predominate (RPT6a, QHK5a-b) while most variants occur at low frequencies. Interestingly, the alleles QHK5a-b and RPT6a are also the two most frequent in an Atlantic salmon population in Northern France (Raynaud et al., 2025). Noting that the aquaculture population genotyped in this study is derived from Northern European populations, this suggests similarities in *PRDM9* diversity over long geographic ranges.

RPT6a and QHK5a-b differ in both ZnF array length and the DNA-contacting residues of the 2nd and 5th ZnFs and are therefore putatively functionally distinct. If these differences translate into divergent binding specificities, the presence of multiple frequent alleles implies a permissive regime for standing functional variation. Such a pattern could arise through genetic drift, reflect an ongoing turnover of *PRDM9* alleles, or represent a stable evolutionary equilibrium. Notably, a recent model predicts that *PRDM9* may experience heterozygote advantage under certain parameter regimes (Z. Baker et al., 2023), which could facilitate the coexistence of functionally divergent alleles at intermediate frequencies.

However, because genotyping in this study was performed in an aquaculture population, founder effects and other non-selective processes associated with artificial breeding may have substantially influenced allele frequencies. Sampling across multiple wild Atlantic salmon populations will therefore be necessary to determine how representative the diversity observed here is of natural populations. Comparisons of diversity patterns across wild populations and across species may further reveal whether similar evolutionary forces act on *PRDM9* in different genomic and demographic contexts. Such analyses will be essential for disentangling demographic effects from selection and for elucidating the evolutionary mechanisms shaping *PRDM9* diversity.

### Dominance interactions between *PRDM9* alleles in Atlantic salmon

Comparisons between linkage maps generated from families where the mother was homozygous or heterozygous for different *PRDM9* alleles revealed a distinct dominance interaction, whereby the alleles QHK5a and QHK5b disproportionately define the recombination localization over RHT10a. A similar suppression of RHT10a by RPT6a may also be present, although this comparison lacked statistical significance. While our study does not resolve the mechanistic basis of these dominance interactions, previous work has shown physical suppression through multimer formation when two PRDM9 variants are present (C. L. Baker, Petkova, et al., 2015) and dominance may be explained by *PRDM9* allele differences in binding site affinity and availability. Over evolutionary time, PRDM9 preferentially erodes its highest-affinity binding sites through biased gene conversion, progressively reducing the availability of effective targets for a given allele (C. L. Baker, Kajita, et al., 2015). Consequently, alleles that have been frequent or long-standing in a population increasingly would be expected to behave recessively when paired with variants whose binding landscapes are less eroded, as hotspot erosion is shown to contribute to dominance interactions between *PRDM9* alleles in mice (Smagulova et al., 2016). In our case, this would imply RHT10a binding sites are more eroded in the European aquaculture progenitor lineage than QHK5a-b/RPT6a. Further work will be required to determine the conditions under which such dominance interactions arise and to identify the molecular and genomic features that correspond to suppression of *PRDM9* allele activity. Such insight will allow for predicting specific allele interactions and the role these interactions play in *PRDM9* evolutionary dynamics.

### Limitations of binding motif inference and screening

In this study, we integrated three complementary analyses that capture recombination at different spatial and temporal scales, together with *PRDM9* sequencing from the corresponding populations. First, a SNP-array based linkage analysis encompassing >100,000 meioses revealed aggregate differences in recombination localization among *PRDM9* genotypes. However, because this analysis relied on low-density SNP data, recombination could only be resolved in 200-kb windows, limiting its utility for identifying PRDM9 binding motifs. To address this constraint, we incorporated two higher-resolution approaches. These included (i) population-averaged recombination estimates resolved at 2-kb windows (Gulbrandsen et al., in prep b) and (ii) a panel of candidate motifs for the alleles RPT6a and QHK5a-b derived from *in silico* binding predictions and *de novo* motif inference from crossover regions in aquaculture families. To evaluate which candidate motifs best reflected PRDM9-directed recombination, we tested their enrichment under recombination hotspots in wild populations with available *PRDM9* genotype data.

The motifs we inferred from crossovers specific to *PRDM9* genotypes may approximate true PRDM9 binding motifs but lack precision. None of the crossover-derived motifs had an enrichment ratio or recombination rate elevation comparable to the motif inferred from population hotspots by Raynaud et al. (2025). This weak signal likely reflects the limited number of informative crossovers available for motif discovery, though it may also be improved by restricting analyses to more informative, PRDM9-linked crossover regions and excluding repeat-rich regions.

In contrast, identifying overrepresented PRDM9-associated sequences from large collections of hotspot regions, such as those derived from WGS-based population-averaged recombination maps, appears to yield more precise binding motifs. However, this approach is inherently biased toward the most frequent alleles in a population and may be confounded when the target regions of multiple *PRDM9* alleles overlap. Because RPT6a predominates across the European Atlantic salmon range, the motif reported by Raynaud et al. (2025) most likely reflects the binding specificity of this variant. While it may be possible to recover motifs for other high-frequency alleles using a similar strategy, this would require populations in which those alleles are abundant. Finally, because population-averaged recombination maps integrate recombination over long evolutionary timescales, historic *PRDM9* allele frequencies and identities may differ from those observed today due to drift or selection. In such cases, present-day hotspot locations may only partially reflect the activity of contemporary *PRDM9* diversity, further complicating motif inference. Continued development of more accurate *in silico* PRDM9 binding prediction tools will likely improve the recovery of current and historically active alleles from population-averaged recombination landscapes.

## Conclusion

In this study, we have shown that the presence of different alleles of *PRDM9* correspond to changes in recombination patterns, confirming PRDM9-directed recombination in Atlantic salmon, as well as demonstrating functional differences between common alleles in Atlantic salmon aquaculture. This further establishes Atlantic salmon as a candidate system for *PRDM9* research in a system removed from mammals, and its implications should be noted for future studies of recombination in Atlantic salmon. We confirmed a previously reported binding motif which likely belongs to the common European allele RPT6a (Raynaud et al., 2025). Future research to determine the target sites of other *PRDM9* alleles and the mechanics underlying dominance interactions will allow us to measure the impacts of different alleles on recombination patterns in Atlantic salmon and expand our understanding of the function and functional diversity of *PRDM9* in this species. Noting an approximate 1.7-million-year divide between North American and European Atlantic salmon populations (Rougemont & Bernatchez, 2018), combined with recent postglacial introgression (Lehnert et al., 2019; Nugent et al., 2024; Rougemont & Bernatchez, 2018), wild Atlantic salmon populations offer an intriguing system to better understand *PRDM9* diversification and evolution over multiple scales.

## Acknowledgements

We would like to thank Bernard de Massy and colleagues at the Université de Montpellier for sharing *PRDM9* primer sequences and helpful discussions. Also, thank you to Tim Knutsen and AquaGen AS for sharing whole genome sequencing data and genotypes and for discussion. The study was supported by PhD financing from the Norwegian University of Life Sciences (NMBU), Faculty of BioSciences (BIOVIT).

## Data availability statement

The authors affirm that all data necessary for confirming the conclusions of the article are present within the article, figures, and tables. *PRDM9* genotyping and allele details are available at https://github.com/Oyvindsg/PRDM9_Atlantic-salmon_aquaculture_genotypes.

## Supplementary

**Figure S1.**
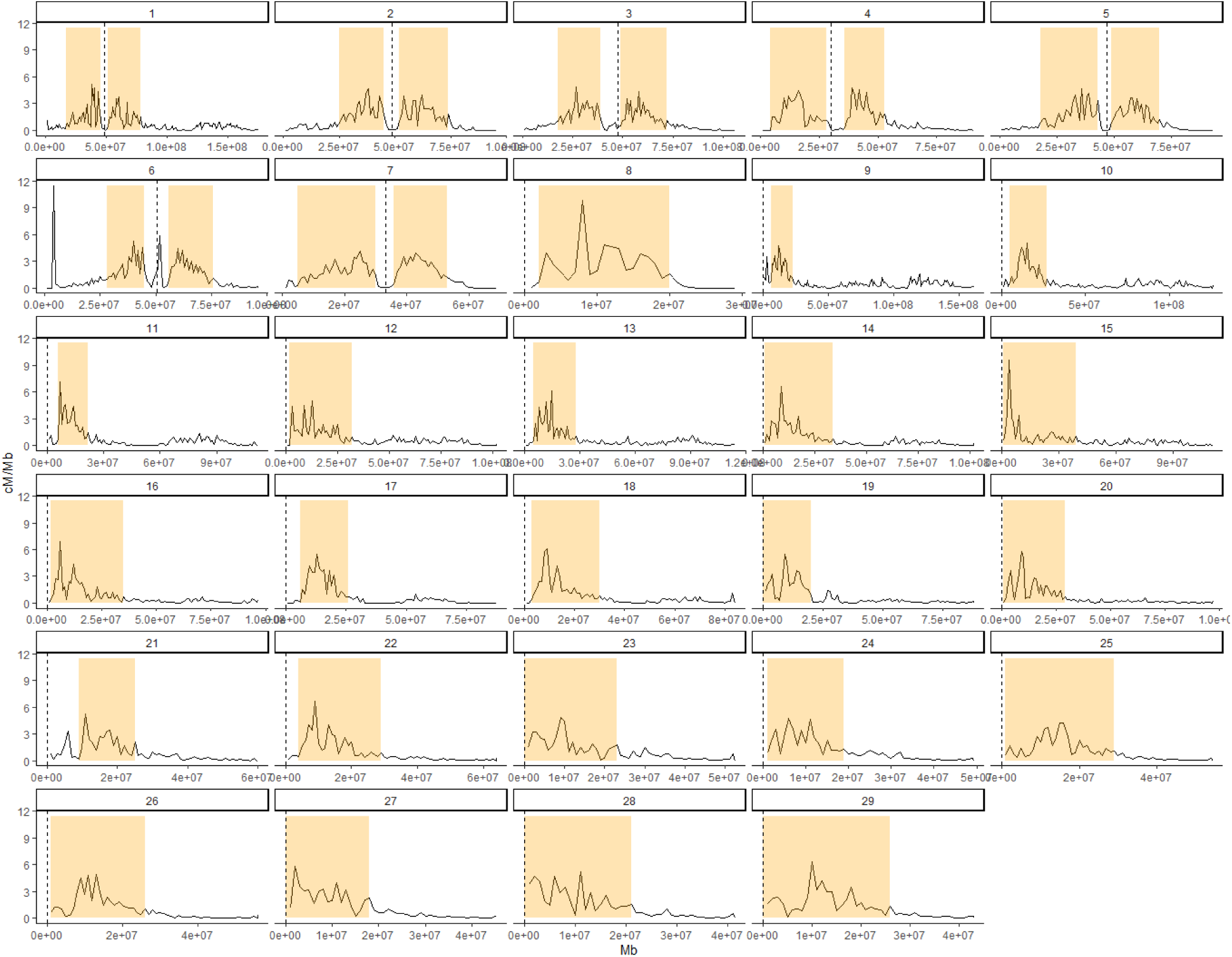
Female Atlantic salmon linkage maps (1 Mb-averaged) across 29 chromosomes. Highlighted regions are recombinationally active and were selected for further analyses. Stapled lines indicate centromere positions.

**Figure S2.**
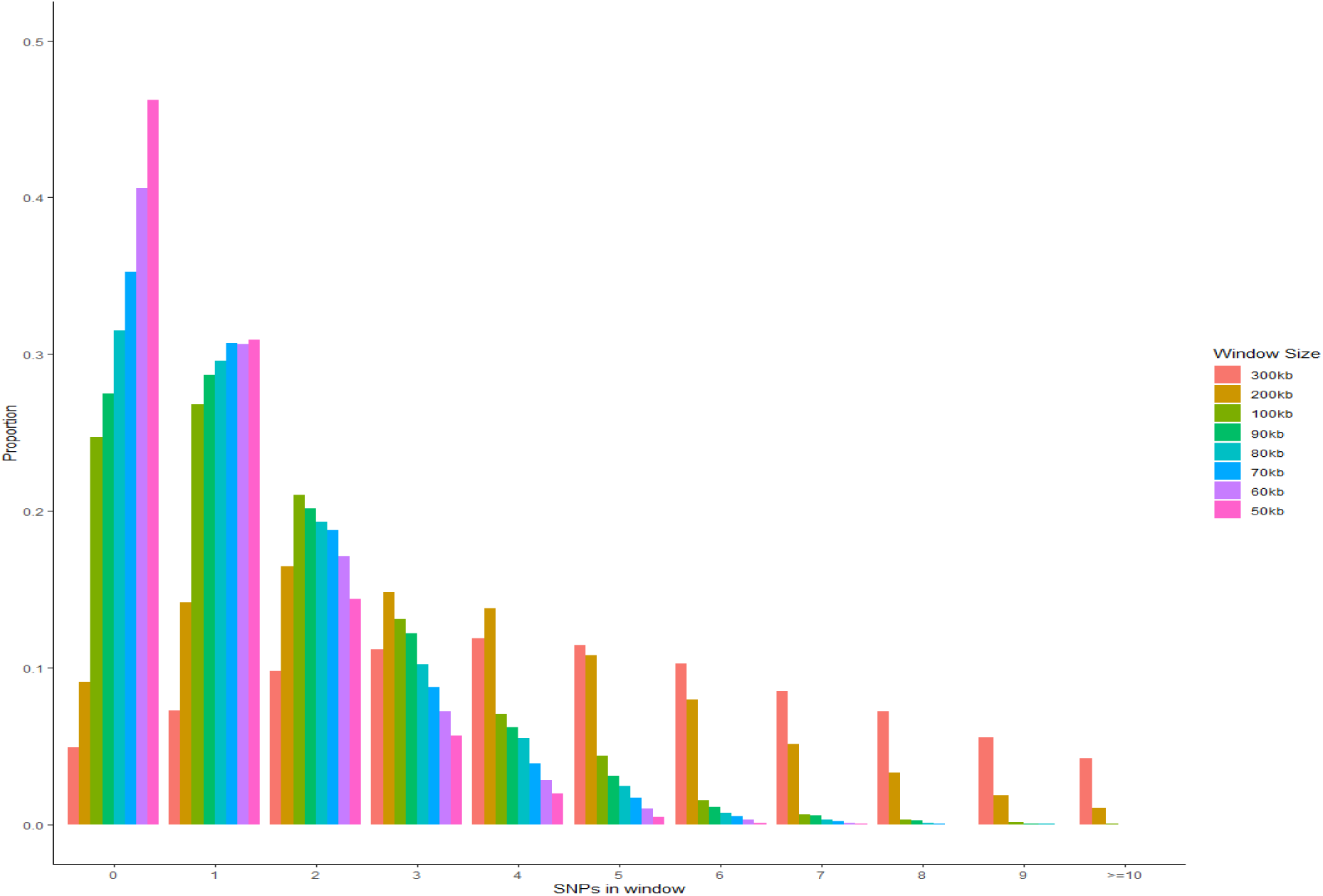
Distribution of the number of SNPs within each window when averaging linkage maps using different window sizes. Window sizes of 400 kb, 500 kb and 1 Mb were also tested but are not shown.

**Figure S3.**
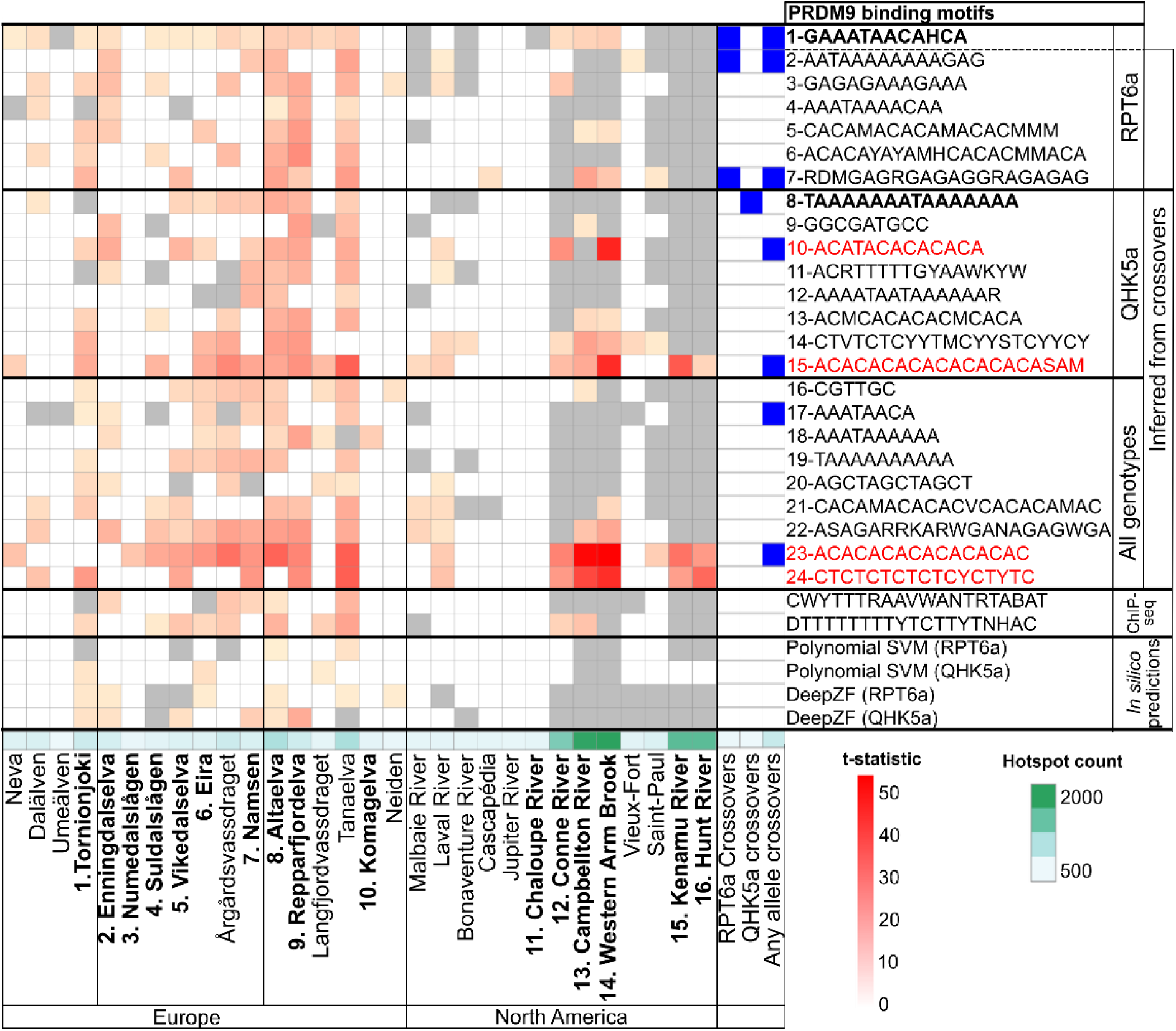
Selected motifs significantly enriched in hotspot regions and their associated increases in recombination intensity in genomic regions containing a PRDM9 binding motif candidate across wild Atlantic salmon populations. The heatmap shows the results of SEA enrichment analyses of hotspot regions, as well as one-sided Welch’s t-tests for recombination rate increases in comparable genomic regions with and without a PRDM9 binding motif candidate (right: motif names, origin and prediction method) across 30 wild Atlantic salmon populations (bottom) with recombination estimates available from Gulbrandsen et al. in prep b). Motifs not significantly enriched in the hotspot set of a population (white) were excluded from recombination rate tests. The increase in recombination at motif-matching sites (t-statistic, yellow to red) is shown for motifs with significant recombination rate increases (Bonferroni-corrected p ≤ 0.05). Motifs significantly enriched in hotspots but without significant recombination rate increases are shown in grey. Crossover regions were not tested for recombination rate increases, but motifs enriched within crossover regions are colored blue. The number of hotspots identified for each population is listed at the bottom of the heat map (white to dark green). Our proposed best binding motif candidates for PRDM9 alleles RPT6a and QHK5a are highlighted in bold and dinucleotide-repeat-like motifs are in red. The ChiP-seq motifs were generated from rainbow trout in Raynaud et al. (2025) and the motif 1-GAAATAACAHCA was inferred from European wild Atlantic salmon population hotspots in the same study.

**Figure S4.**
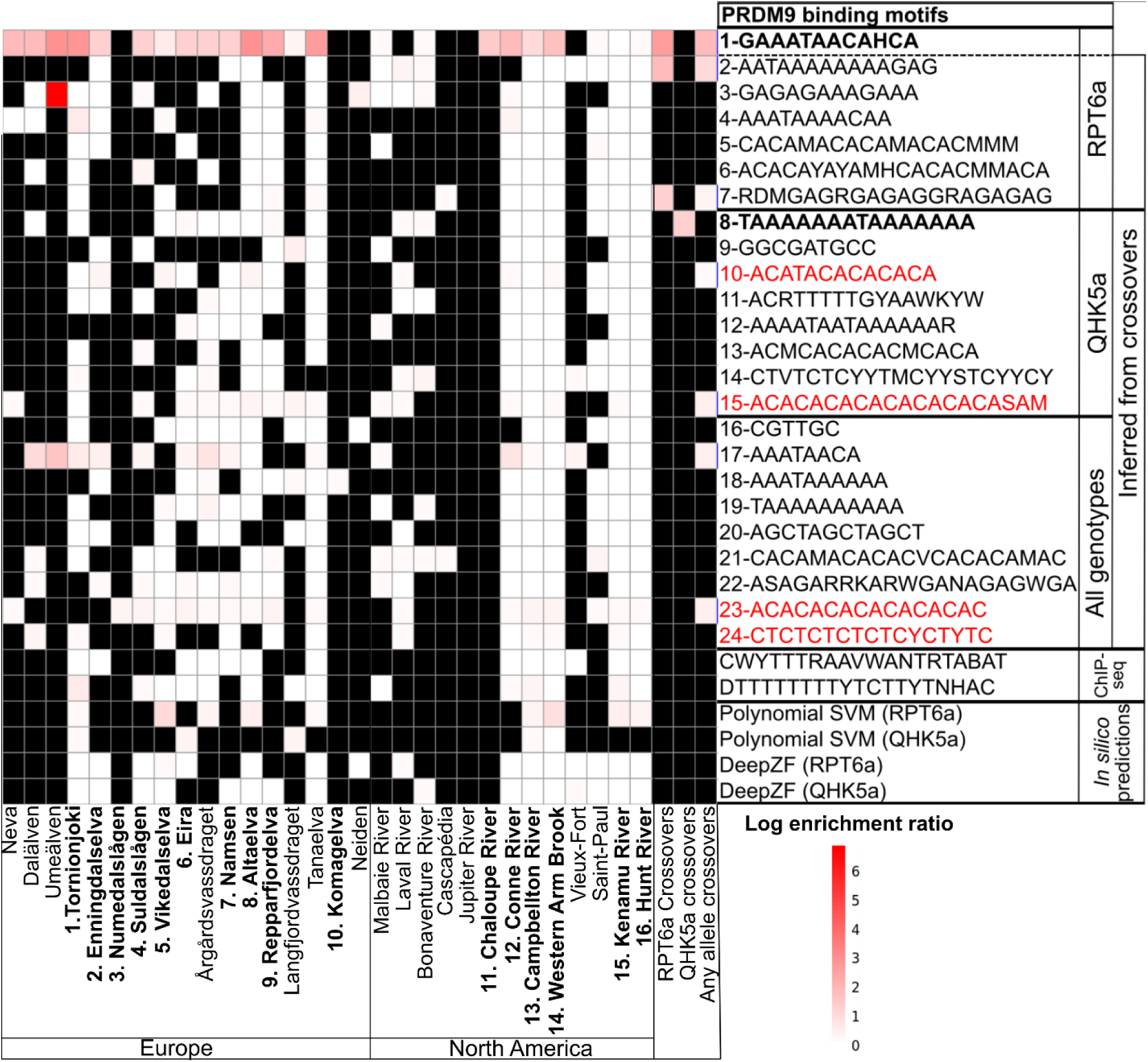
Log-scale enrichment ratios (light yellow to red) for motifs across wild Atlantic salmon populations. Motifs not found by SEA to be enriched in hotspots are in black. Refer to Figure S3 for explanations of motifs and populations.

**Figure S5.**
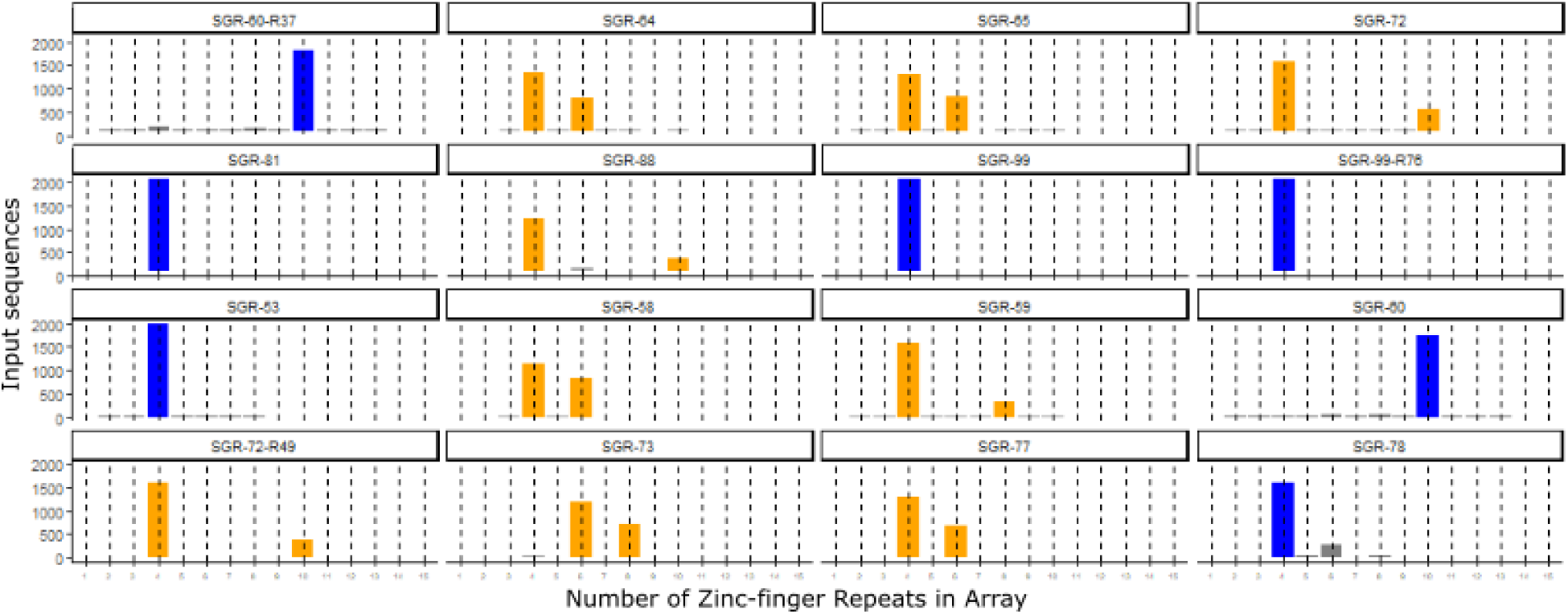
Lengths of alleles accepted for genotyping using modified rules. Colored bars show the lengths of the alleles passing genotype reporting thresholds. Blue = Homozygous genotype, Orange = Heterozygous genotype, Grey = Sequences failed to pass genotyping.

**Figure S6.**
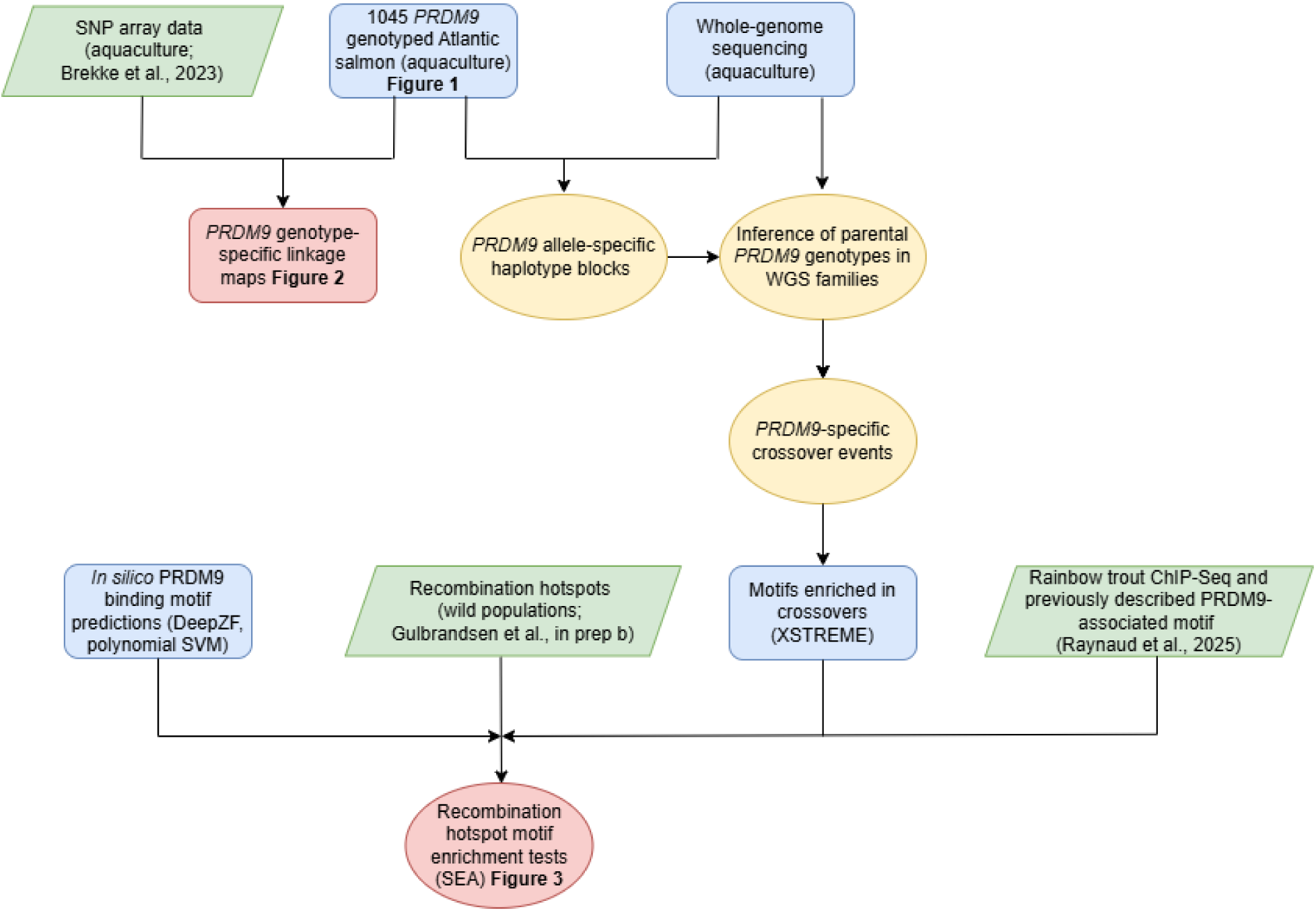
Overview of datasets, methods and analyses utilized in this study.

**Table S1.**
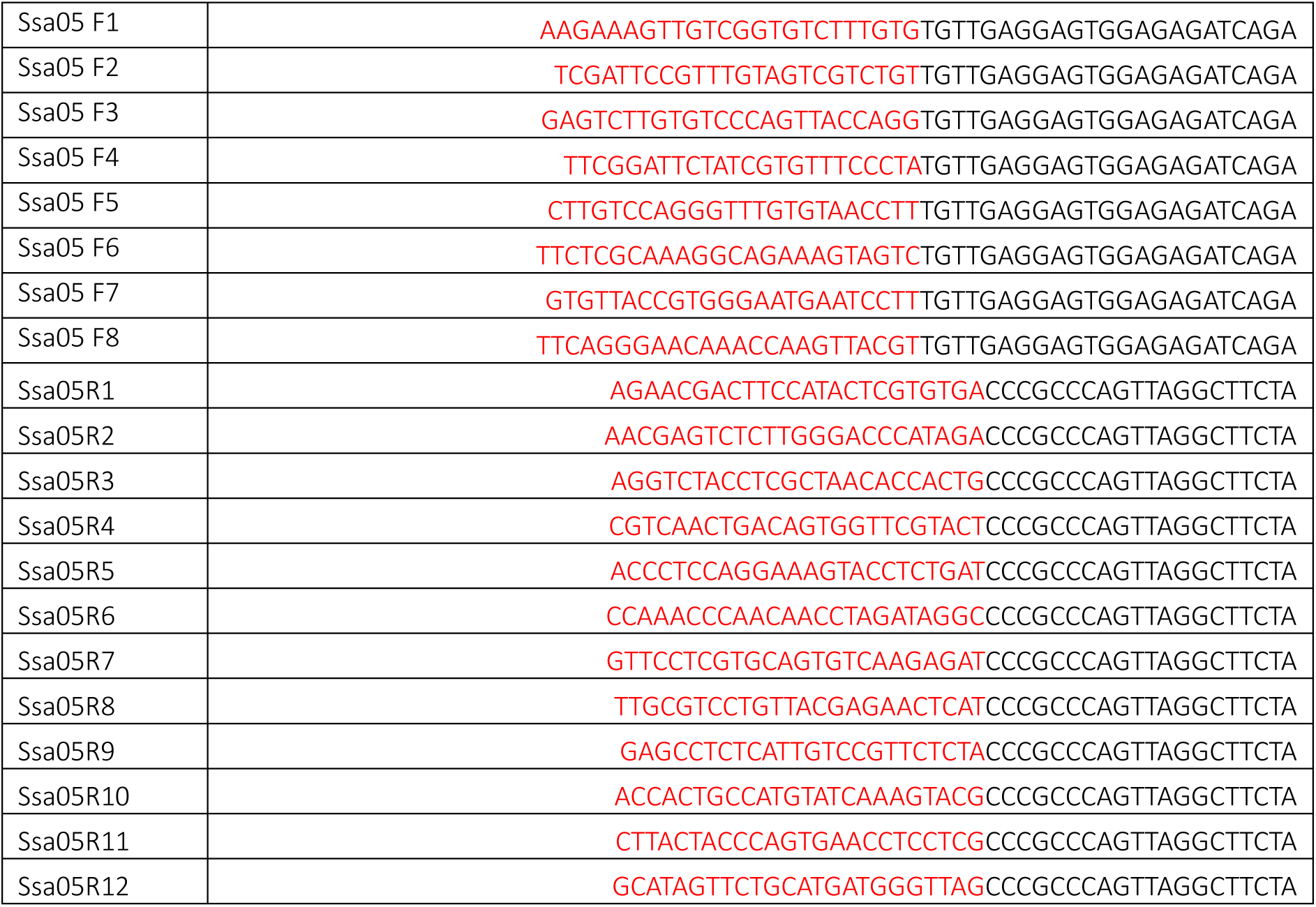
Oligo sequence (5’ to 3’) of primers, including PRDM9 annealing sequence (black) and unique barcodes (red).

**Table S2.**
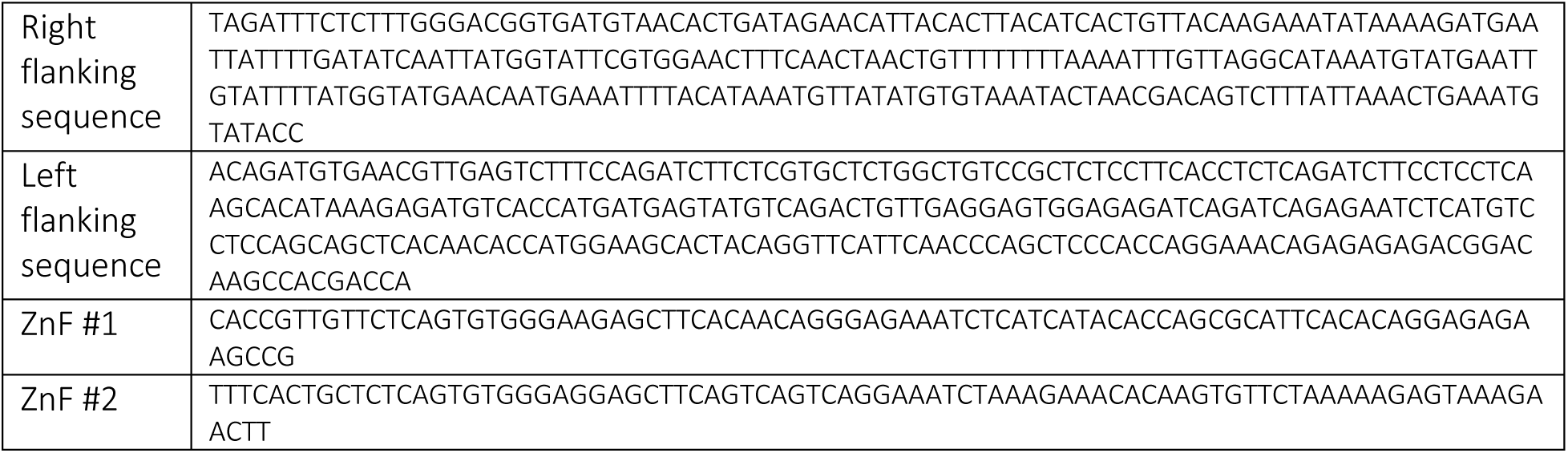
Atlantic salmon sequence substitutes for sequences used in the PRDM9 genotyper developed by Alleva et al. (2021).

**Table S3.** Ratios of different-length haplotypes utilized when determining PRDM9 genotype. Table is taken from Alleva et al (2021), with our modified ratios in bold (rules 3 and 4). Rules are explained as follows (description from Alleva et al., 2021, ZF = Zinc Finger): “Where fi, fj, and fk are the frequencies of the most frequent (i), second most frequent (j), and third most frequent (k) ZF arrays. Rules are processed consecutively; thus, an individual where the ZF array lengths can be inferred using rule 1 will not be tested by further rules.”

| # | Rule | Length (hap1) | Length (hap2) |
| --- | --- | --- | --- |
| 1 | All zinc finger arrays had $i$ zinc fingers ($f_i = 1$) | $i$ | $i$ |
| 2 | $f_i + f_j \geq 0.7$ AND $f_i / f_j < 2$ | $i$ | $j$ |
| 3 | $f_i + f_j \geq 0.7$ AND $f_i / f_j < 5$ AND $f_j / f_k > 3$ | $i$ | $j$ |
| 4 | $f_i + f_j \geq 0.7$ AND $f_i / f_j > 5$ | $i$ | $i$ |
| 5 | $f_i \geq 0.7$ | $i$ | $i$ |

**Table S4.** Number of occurrences of phased haplotypes in the linkage block surrounding PRDM9 in the presence or absence of specific PRDM9 alleles. Only haplotypes which exclusively co-occurred with a single allele were used for imputation. Only parents homozygous for PRDM9 alleles RPT6a or QHK5a after imputation were used for allele specific crossover sets.

| Allele | Haplotype | # With allele and haplotype | # with allele, without haplotype | # with haplotype, without allele | # without haplotype and allele |
| --- | --- | --- | --- | --- | --- |
| RPT6a | 1 | 4 | 80 | 0 | 22 |
| RPT6a | 2 | 41 | 43 | 0 | 22 |
| RPT6a | 3 | 6 | 78 | 0 | 22 |
| QHK5a | 4 | 6 | 40 | 0 | 58 |
| QHK5a | 5 | 17 | 29 | 0 | 58 |
| QHK5b | 6 | 5 | 5 | 0 | 96 |
| RHT10e | 7 | 3 | 7 | 0 | 96 |
| RPT6e | 8 | 2 | 2 | 0 | 102 |
| QHK8b | 9 | 2 | 2 | 0 | 102 |
| QDT9a | 10 | 6 | 8 | 0 | 92 |
| QDT6b | 11 | 1 | 1 | 0 | 104 |

